# Degradation of Lon in *Caulobacter crescentus*

**DOI:** 10.1101/2020.06.12.149492

**Authors:** Benjamin B Barros, Samar A Mahmoud, Peter Chien, Rilee D. Zeinert

## Abstract

Protein degradation is an essential process in all organisms. This process is irreversible and energetically costly; therefore, protein destruction must be tightly controlled. While environmental stresses often lead to upregulation of proteases at the transcriptional level, little is known about post-translational control of these critical machines. In this study we show that in *Caulobacter crescentus* levels of the Lon protease are controlled through proteolysis. Lon turnover requires active Lon and ClpAP proteases. We show that specific determinants dictate Lon stability with a key carboxy-terminal histidine residue driving recognition. Expression of stabilized Lon variants results in toxic levels of protease that deplete normal Lon substrates such as the replication initiator DnaA to lethally low levels. Taken together, this work demonstrates a feedback mechanism in which ClpAP and Lon collaborate to tune Lon proteolytic capacity for the cell.

**Importance:** Proteases are essential, but unrestrained activity can also kill cells by degrading essential proteins. The quality control protease Lon must degrade many misfolded and native substrates. We show that Lon is itself controlled through proteolysis and that bypassing this control results in toxic consequences for the cell.

## Introduction

Energy dependent protein degradation in bacteria is carried out by the conserved <u>A</u>TPase <u>A</u>ssociated with diverse cellular <u>A</u>ctivities (AAA+) proteases Lon, FtsH, HslUV, ClpAP, and ClpXP proteases (1, 2). The protease must first recognize a motif, known as a degron, in the tertiary or primary amino acid sequence to begin the terminal destruction of targets. Once the degron is recognized, the substrate is mechanically unfolded using the power of ATP hydrolysis and then translocated into the proteolytic chamber where it is destroyed. In the case of ClpAP and ClpXP, two separate polypeptides that encode an unfoldase (ClpA and ClpX) and a peptidase (ClpP) come together to form the active protease complex. By contrast, the bacterial Lon protease is encoded on a single polypeptide that contains both an unfoldase and peptidase domain.

Transcriptional regulation by heat stress-induced alternative sigma factors is a well understood mechanism used by bacteria to modulate the heat shock response (3, 4). During the bacterial heat shock response, accumulation of unfolded substrates is sensed by the DnaK/RpoH system and results in increased transcription from genes with heat shock promoter elements (5–7). Unfolded substrates arising from heat stress must either be refolded or destroyed, therefore it is not surprising that chaperones and proteases are the primary components of the RpoH driven heat-shock induced regulon (3). In this way, protein homeostasis factors are appropriately upregulated during conditions when the demand is high. Post-translational control of chaperones/proteases are poorly understood but given that these stress responses are both universal and integral to life, it seems likely that more complex regulation is at play.

In this study, we find that in *Caulobacter crescentus*, Lon levels are regulated through proteolysis that requires active Lon and ClpAP proteases. Stability of Lon depends on exposure of its native C-terminus, which contains a key histidine needed for Lon degradation. *In vitro* assays suggest an additional component beyond Lon and ClpAP is needed for degradation of Lon. Expression of nondegradable variants of Lon results in loss of viability and aberrant morphology. This toxic effect is likely due to a drastic reduction in levels of essential Lon substrates, such as the replication initiator DnaA, when Lon is both stabilized and expression is increased. This discovery that Lon is regulated via its own degradation supports models where levels of proteases must be tightly controlled in order to sustain normal growth, while ensuring sufficient capacity to handle stressful conditions.

## Results

### Lon degradation *in vivo* requires Lon catalytic activity

Monitoring protein levels during translational inhibition is a well-established proxy for protein degradation. During the course of our studies exploring the degradation of the Lon substrate DnaA, we were surprised to observe the loss of Lon as well during these assays (**Figure 1**). By contrast, levels of the ClpP protease were constant throughout the time course of our experiment. These data suggest that Lon levels are controlled by degradation in *Caulobacter crescentus*.

**Figure 1.**
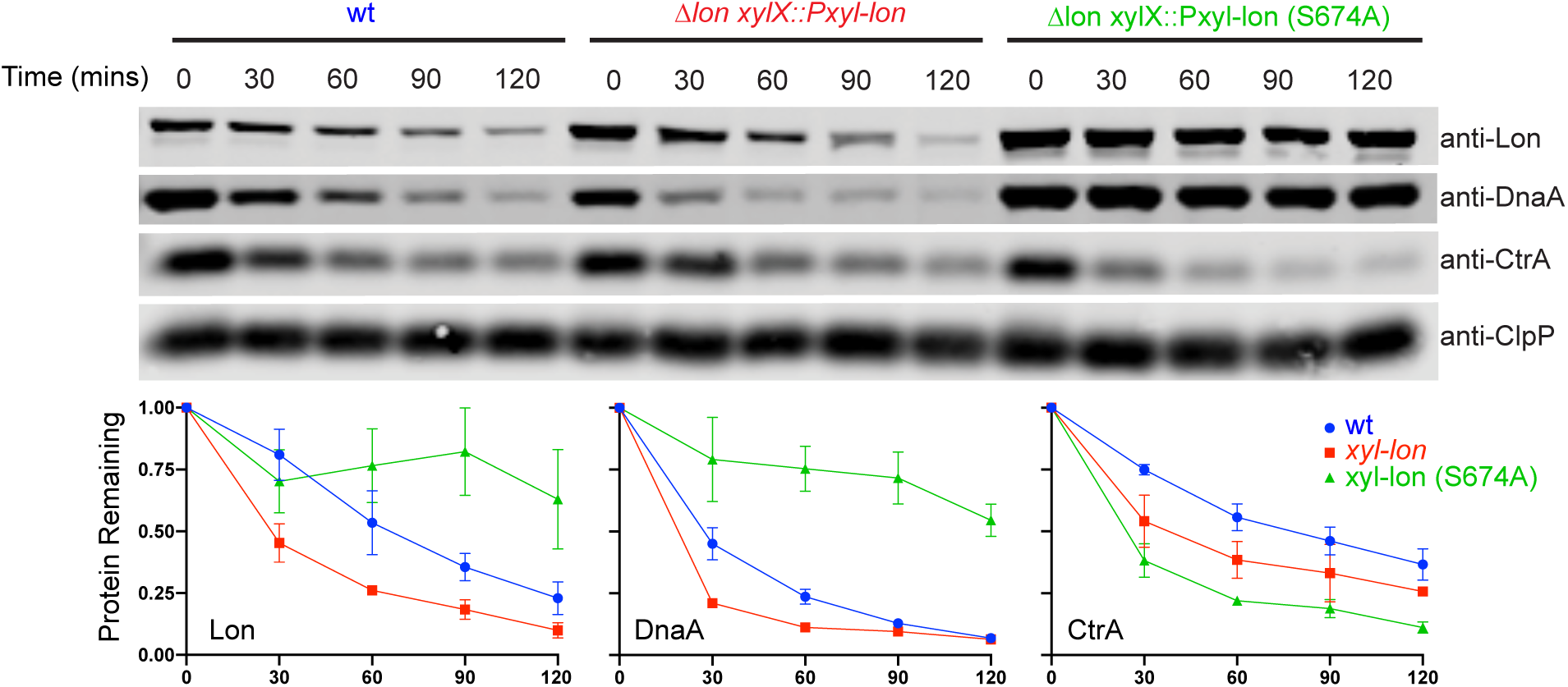
Lon degradation *in vivo* requires Lon catalytic activity. *In vivo* degradation of Lon, CtrA, DnaA and ClpP following translation shutoff in wt, Δ*lon xylX::lon*, Δ*lon xylX::lon* S674A. Quantification of biological triplicates shown below.

We next asked if Lon was responsible for its own degradation *in vivo*. To test this, we made strains where the wild-type (wt) or a catalytically inactive (S674A) allele of Lon was integrated at the inducible *xylX* locus in a Lon-deficient strain. As we observed for the endogenous Lon, induced Lon was robustly degraded following translational shutoff, but interestingly, the turnover of the catalytically inactive Lon S674A was greatly reduced (**Figure 1**). As expected, levels of the Lon substrate DnaA remained stable in this background (**Figure 1**). To ensure that the maintenance of Lon levels in this background was due to loss of degradation and not a failure of translation inhibition, we measured levels of CtrA, a ClpXP-dependent substrate (8). As expected, CtrA was degraded regardless of Lon background, supporting our conclusion that Lon degradation requires intact Lon catalytic activity.

### The C terminus of Lon is necessary for degradation

Degrons are frequently located at the exposed termini of protein substrates (1). We asked whether the exposed N or C terminus of Lon was required for recognition by appending an M2-FLAG epitope to either terminus and integrating these constructs at the inducible *xylX* locus in both wildtype and Δ*lon* strains. Following induction, steady state levels of the C-terminally tagged Lon (lonM2) were substantially higher than the other constructs in both wildtype and Δ*lon* (**Figure 2 A**). Consistent with this, induction of all Lon variants in the Δ*lon* background resulted in loss of DnaA, but this decrease was most pronounced when lonM2 was expressed (**Figure 2 A**).

**Figure 2.**
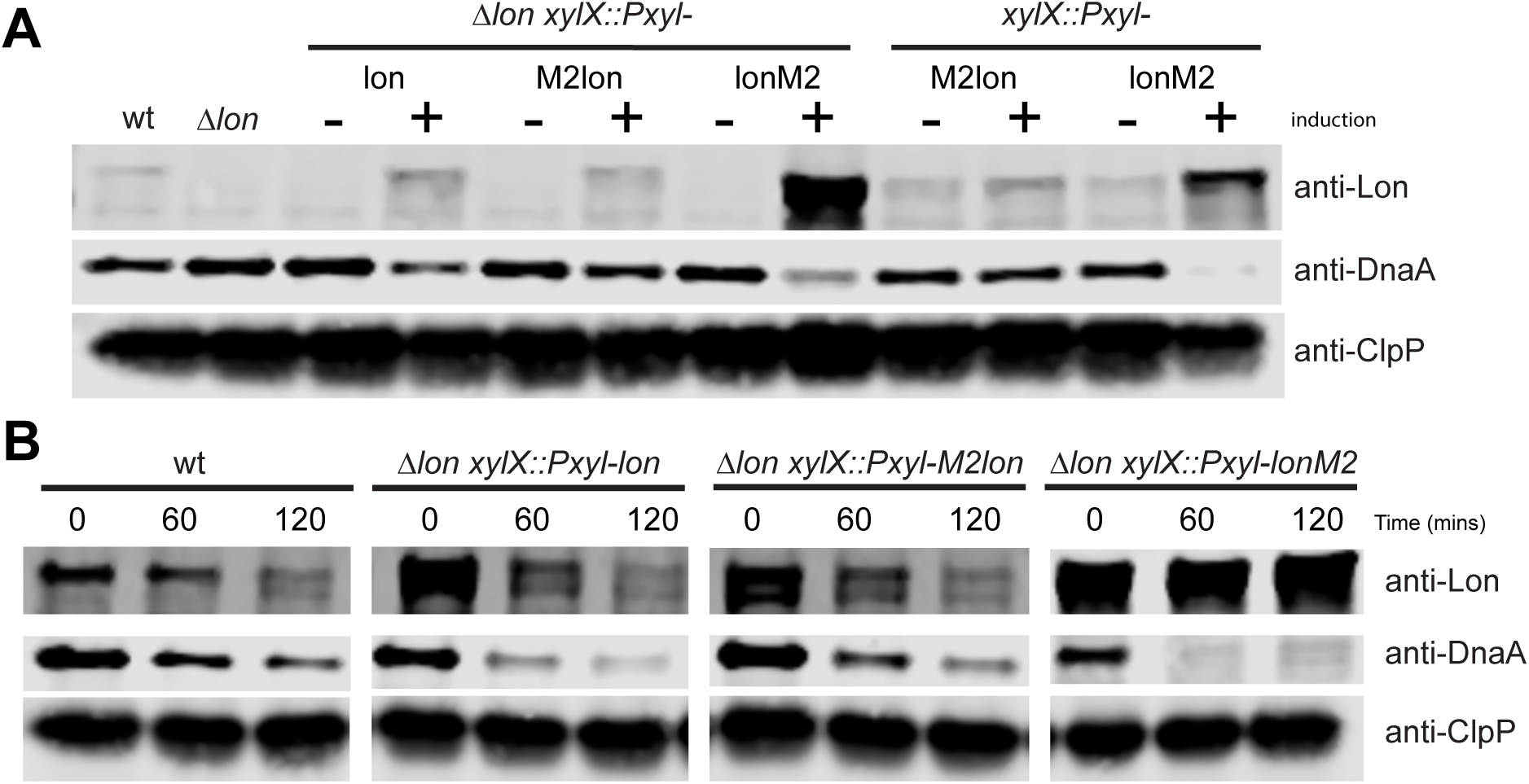
The C terminus of Lon is necessary for degradation. (A) Steady state protein levels of Lon, DnaA and ClpP in wt or Δ*lon* strains alone or with Lon, amino terminus M2-FLAG epitope Lon (M2lon), or carboxy terminus M2-FLAG epitope Lon (lonM2). Samples were normalized to starting OD_600_ in lysis buffer prior to Western Blot analysis. Cropped images of western blots probing for Lon, DnaA, and ClpP. (B) *In vivo* stability of Lon following translation inhibition in wt, Δ*lon xylX::lon*, Δ*lon xylX::M2lon*, Δ*lon xylX::lonM2*. Levels of DnaA are shown as a control for Lon proteolytic activity and ClpP as a loading control.

We next tested whether the increase in steady-state levels of Lon was reflected by changes in protein stability. While the untagged (Lon) and N-terminally tagged (M2Lon) were clearly degraded upon translational shutoff, the C-terminal tagged (LonM2) was stable during our timecourse (**Figure 2 B**). As expected, the rate of DnaA degradation correlated well with the observed changes in Lon levels. Taken together with our prior observations, our working model is that the C-terminus of Lon contains a degron that facilitates Lon degradation in a manner that requires active Lon protease.

### Overexpression of a non-degradable Lon results in toxicity and aberrant morphology

In *E. coli*, overproduction of Lon results in lethality due to degradation of a bacterial anti-toxin and subsequent buildup of the corresponding toxin protein (9). However, when we overexpressed native Lon in *Caulobacter*, we saw little phenotypic consequence (**Figure 3 A**). Interestingly, expression of the stabilized lonM2 construct resulted in reduced viability on solid agar media that was especially apparent in cells carrying both a native copy of *lon* and expressed the lonM2 construct (**Figure 3 A**). Microscopic examination of induced cells showed that expression of lonM2 resulted in filamentous cells, while expression of either the wildtype or M2lon variant did not (**Figure 3 B**). These filamentous cells were smooth and markedly straighter as measured by quantification of curvature (**Figure 3 C**). Thus, the inability to degrade Lon makes cells particularly sensitive to excess amounts of this protease.

**Figure 3.**
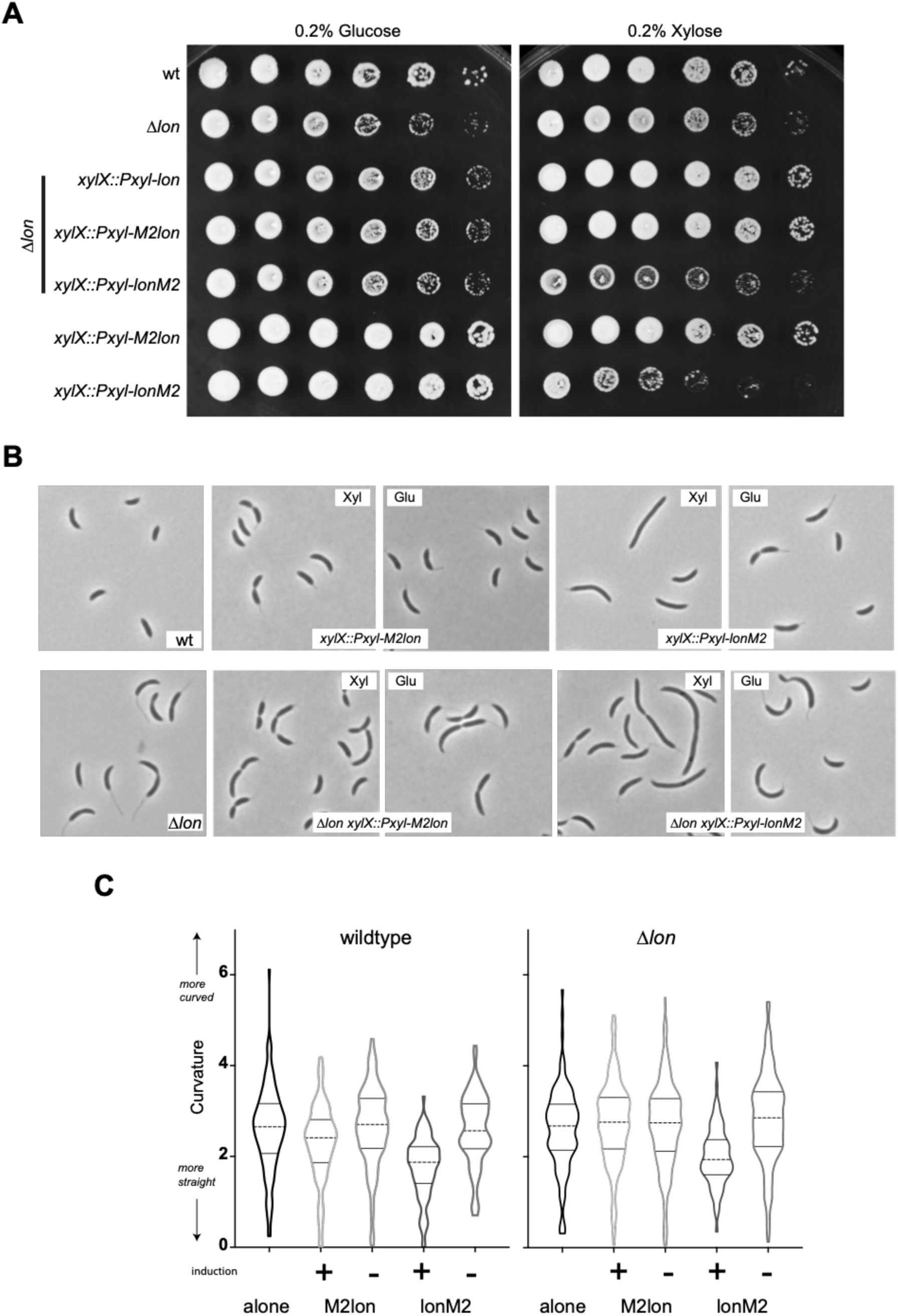
Overexpression of a non-degradable Lon results in toxicity and aberrant morphology. (A) Induction of *lonM2* from the *xylX* promoter results in cell death in wt and the Δlon strain. Cells were grown to exponential phase in 0.2% glucose, normalized by OD_600_, serially diluted 10-fold and spotted onto media supplemented with 0.2% glucose (-induction) or 0.2% xylose (+ induction). Images of plates were taken on day 2 of growth. (B) Induction of lonM2 from the *xylX* promoter results in aberrant morphologies of wt and Δ*lon xylX::M2lon or xylX::lonM2*. Representative images of indicated strains grown in exponential phase in the presence of 0.2% xylose or 0.2% glucose for 6 hours prior to imaging. Cell curvatures (n>100) were quantified using microbeJ and graphed with Prism.

### The C-terminal histidine is necessary for degradation of Lon

Given that the C-terminus of Lon appeared critical, we performed a multiple sequence alignment to compare Lon across multiple bacteria species. We found that *Caulobacter* Lon encodes a slightly longer C-terminal extension that ends with a histidine residue, an organization conserved in *α*-proteobacterium. More distant species such as *Escherichia coli* and *Bacillus subtilis* lack the C-terminal extension and do not encode histidine as the terminal residue (**Figure 4 A**). Interestingly, the *E. coli* Lon substrate SulA contains a C-terminal histidine shown to be crucial for its degradation (10), so we next tested if this same motif is important for *Caulobacter* Lon degradation.

**Figure 4.**
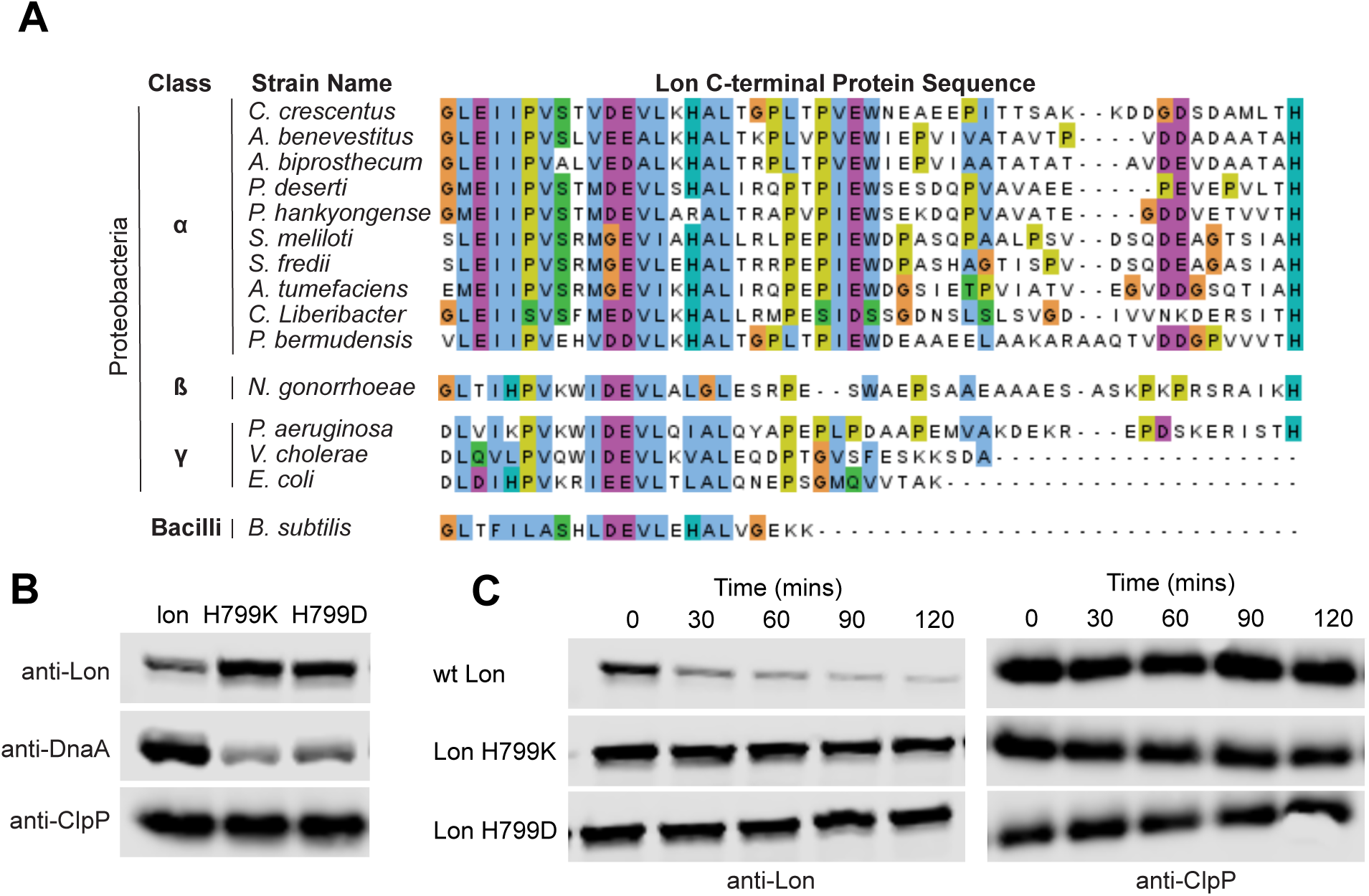
A C terminal histidine on Lon is necessary for stability. (A) Structural alignment of the C-terminus of Lon. (B) Steady state protein levels of Lon DnaA and ClpP in Δ*lon xylX::lon* (lon native, lon H799K, lon H799D) strains. Samples were normalized to starting OD_600_ in lysis buffer prior to Western Blot analysis. Cropped images of western blots probing for Lon, DnaA, and ClpP. (C) *In vivo* stability of Lon following translation inhibition in Δ*lon xylX::lon* (wildtype lon, lon H799K, lon H799D) strains. Levels of DnaA are shown as a control for Lon proteolytic activity and ClpP as a loading control.

We mutated residue H799 to a lysine or aspartic acid (H799K or H799D), expressed these constructs from the *xylX* promoter, and monitored levels and stability of these Lon variants. Steady state levels of Lon H799K and H799D were significantly higher than the wildtype allele and, consistent with them being functional, DnaA levels were dramatically reduced (**Figure 4 B**). To monitor the stability of the C-terminal mutants, Lon levels were measured after inhibiting protein translation. Mutation of the C-terminal histidine residue to a Lysine or Aspartic acid residue (LonH799K and LonH799D) resulted in a dramatic stabilization of Lon (**Figure 4 C**). We conclude that the terminal histidine residue of *Caulobacter* Lon is a critical element of the Lon degron.

### Degradation of Lon requires additional factors including ClpA

We next tested whether Lon was degrading itself, as our previous results showed that Lon activity was needed for Lon degradation *in vivo* (**Figure 1**). We purified *Caulobacter* Lon, but did not find any sign of significant autodegradation, even though the known Lon substrate DnaA was degraded well in these assays (**Figure S1**). This implied that another factor might be needed *in vivo* for robust degradation.

In *Caulobacter crescentus*, ClpAP was identified as an auxiliary protease that also degrades the Lon substrate DnaA (11). DnaA activity is essential for replication of initiation and as a transcription factor (12) which implies that its levels would be critical for viability. We hypothesized that ClpAP could provide a logical feedback loop, coupling Lon levels with ClpAP activity to tune the amount of degradation of DnaA. Interestingly, deletion of *clpA* resulted in a significant increase in the steady state levels of Lon and a reduction in known Lon substrates DnaA and CcrM (**Figure 5 A**). Translational shutoff experiments showed that Lon degradation was reduced in the Δ*clpA* strain, supporting a possible role for ClpA or ClpAP in degrading Lon (**Figure 5 B**). However, adding ClpAP to our *in vitro* reactions was insufficient to degrade Lon (**Figure S1**). The failure of Lon to be degraded in these reduced systems indicates either that our *in vitro* system does not sufficiently mimic *in vivo* conditions or that additional factors are needed that connect Lon and ClpAP activities.

**Figure 5.**
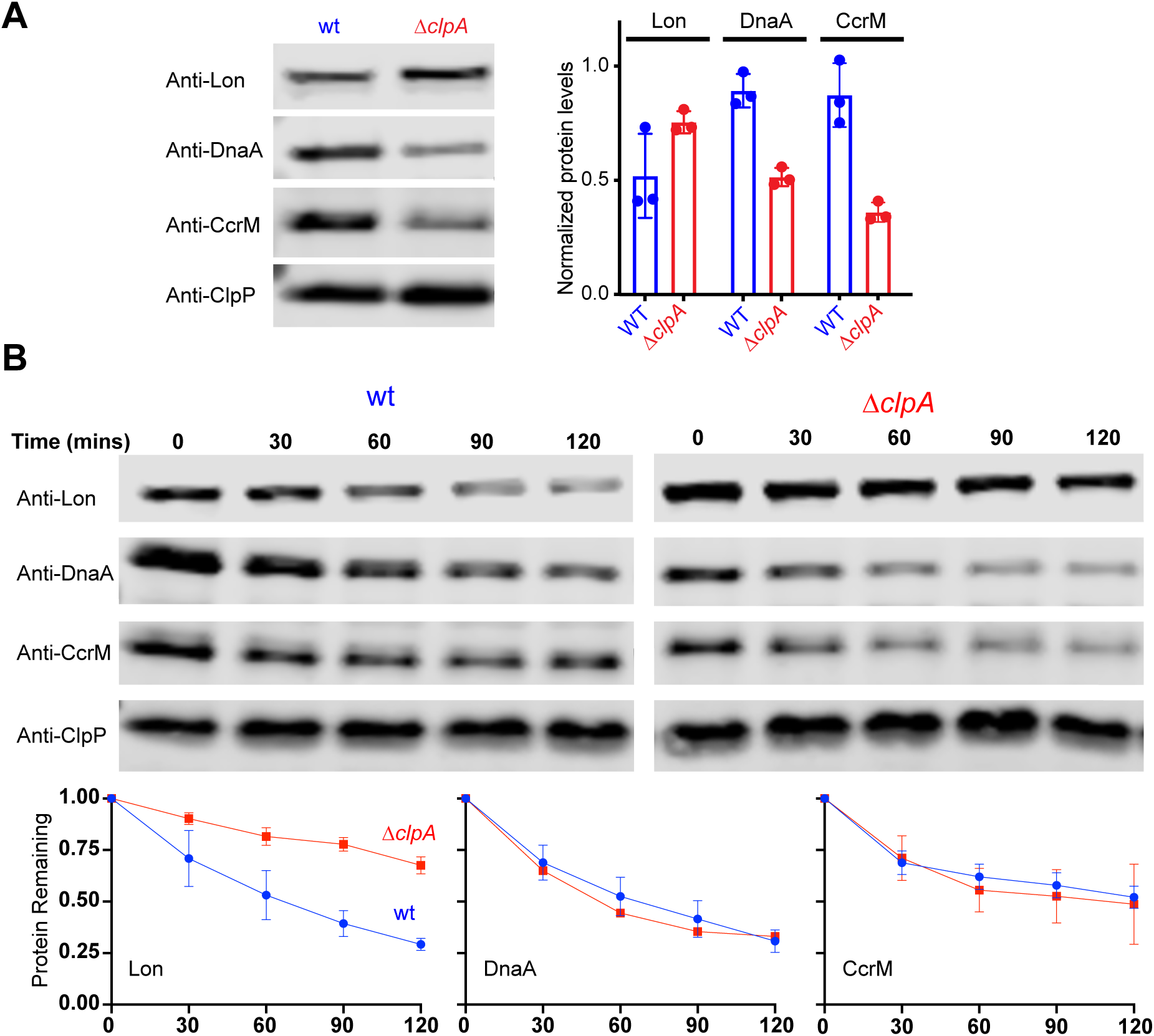
*in vivo* degradation of Lon is dependent on ClpA. (A) Steady state protein levels in wt and *clpA* deletion strain. Representative images of western blots using anti-Lon, DnaA, CcrM. Anti-ClpP was used as a loading control. Protein quantifications of biological triplicates with error bars represented as standard deviation. (B) Protein stability of Lon in wt and Δ*clpA* strains. Strains were grown to exponential phase at 30°C prior to inhibition of protein synthesis by addition of chloramphenicol. Samples were withdrawn at the indicated time points, normalized to starting OD_600_ in lysis buffer. Lysates were used for Western blot analysis and probed with antibodies to Lon, DnaA, CcrM and ClpP. Cropped representative images of biological triplicates with quantifications of biological triplicates are shown with error bars represented by standard deviation.

## Discussion

Energy-dependent proteases help establish steady-state levels of proteins *in vivo (1)*. Because proteolysis is irreversible, the levels of proteases must be highly controlled to meet the demands of the cell in ensuring steady state levels while maintaining the ability to respond to stress. While other protease components have been shown to be degraded, such as the case for ClpA autodegradation by ClpAP (13, 14), the biological impact of this degradation has not been well explored.

In this study we show that in *Caulobacter*, the Lon protease is controlled post-translationally through degradation (**Figure 6**). The instability of Lon *in vivo* requires active Lon and ClpAP proteases suggesting a molecular link between these two pathways. However, we could not reconstitute Lon degradation *in vitro* with just Lon and ClpAP. Therefore, we hypothesize that an additional regulator Y (**Figure 6**) is needed to facilitate Lon degradation by either ClpAP or Lon itself. In this model, either Lon or ClpAP would degrade X, which is an inhibitor of Y. This would then make proteolysis of Lon dependent on activity of both Lon and ClpAP, and would explain why we are unable to recapitulate this behavior with only these purified proteins.

**Figure 6.**
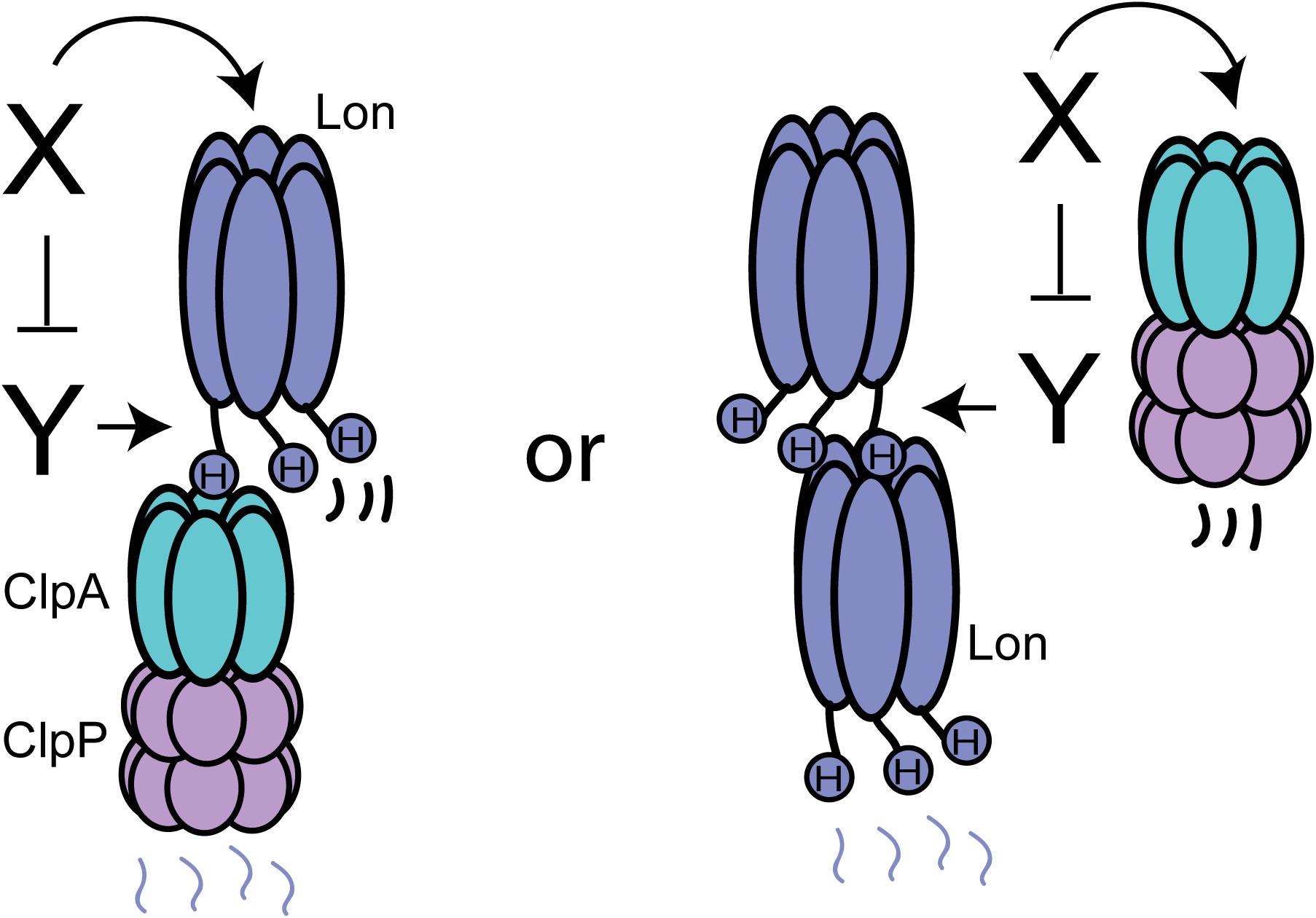
Lon stability is dependent on ClpAP and Lon activity. Lon degradation is dependent on ClpAP and Lon activity, but these proteases are insufficient on their own *in vitro*. This suggests a need for a factor (Y) that enhances degradation of Lon dependent on a C-terminal histidine residue by either ClpAP or Lon itself. An inhibitor (X) of Y is degraded by either Lon or ClpAP, thereby linking the in vivo observations with our *in vitro* biochemistry.

Degradation of Lon requires a free C-terminal histidine (**Figure 4 A; Figure 6**) that is conserved in *α*-proteobacteria and is also found in some other bacterial species. It would be fascinating to determine if Lon is also unstable in other species containing the conserved C-terminal histidine, such as the pathogen *Pseudomonas aeruginosa* (**Figure 4 A**). In *Caulobacter*, we hypothesize that the toxic consequence of expressing stabilized Lon is due to loss of an essential substrate, likely DnaA, given that cells depleted of DnaA develop a filamentous morphology (15) similar to those we see with expression of lonM2 (**Figure 3 B**). However, it is possible that excess antitoxin degradation such as seen in the case of Lon overexpression in *E coli* (9) also contributes to the loss in viability.

The overexpression of a protein is an artificial tool that can be used to test the most extreme aspects of a model. We show here that overexpression of a non-degradable Lon resulted in cell death (**Figure 3 A**). However, this begs the question of whether there are conditions in which Lon protease degradation is required when it is under endogenous transcriptional control? We reason that Lon levels would be particularly important during heat shock conditions where the Lon protease has been shown to rapidly degrade DnaA in the presence of unfolded substrates thereby arresting growth (16). This growth arrest protects cells during extreme thermal stress conditions, but they must eventually resume growth when the damage subsides. Because Lon degrades essential factors necessary for DNA replication and division such as DnaA (16) and CcrM (17) it is easy to envision that recovery from these conditions might be delayed if sustained elevated levels of Lon prevent the accumulation of these substrates. Additionally, competition between substrates for a limited pool of protease has been recently highlighted as a mechanism for toggling growth under stressful conditions (18). Therefore, cells likely have a need to maintain low levels of proteases during certain growth or stress responses, which can be facilitated by post-translational controls such as what we have shown here.

## Methods

### Bacterial strains, media and growth conditions

*Caulobacter crescentus* NA1000 strains were grown in peptone (2 g/L) yeast (1 g/L) extract medium (PYE) supplemented with 1 mM MgSO_4_, 0.5 mM CaCl_2_ at 30°C with shaking at 200 RPM. To generate strains, antibiotics were added to PYE supplemented with 1.5% agar at the final concentrations: kanamycin (25 µg/ml), gentamycin (5 µg/ml), spectinomycin (100 µg/ml), and oxytetracycline (2 µg/ml). After the initial selection steps of strain generation, antibiotics were excluded from all liquid and solid growth assays. For induction purposes, the xylose promoter was induced with xylose (0.2%) or repressed with glucose (0.2%). *E. coli* strains were grown in Luria-Bertani broth (LB) at 37°C with shaking at 180 RPM. The cultures were supplemented with the following antibiotic concentrations: oxytetracycline (15 µg/ml), kanamycin (50 µg/ml), gentamycin (20 µg/ml), spectinomycin (100 µg/ml), chloramphenicol (30 µg/ml), or ampicillin (100 µg/ml). The list of strains and plasmids are provided in Table S1 in the supplemental materials.

### Strain construction

All *Caulobacter* strains were derivatives of CPC176, an isolate of strain CB15N/NA1000.

For overexpression or merodiploids, strains were constructed using a one-step recombination procedure at the xylose locus using pXGFPN-5 or at the endogenous locus, using pMCS-1 (19).

CPC816 was constructed by electroporating plasmid pXGFPN-5_CcLon (S674A) into CPC609 and selecting onto oxytetracycline plates.

CPC920 and CPC922 were generated by electroporating pXGFPN-5_Cclon(M2) into CPC176 and CPC609 with selection on oxytetracycline plates.

CPC921 and CPC923 were generated by electroporating pXGFPN-5_(M2)Cclon into CPC176 and CPC609 with selection on oxytetracycline plates.

CPC929 and CPC930 were generated by electroporating pXGFPN-5_CclonH799K or pXGPFN-5_CclonH799D respectively into CPC609 with selection on oxytetracycline plates.

CPC816, CPC920, CPC921, CPC922, CPC923, CPC924, CPC 929, and CPC930 were all validated by anti-Lon and anti-flag (when appropriate) western for clones that expressed Lon with xylose and had low levels of Lon with no induction.

### Plasmid Construction

#### Integration plasmids

All Lon related constructs for integration at the xylose locus were generated using the vector pXGFPN-5. For native constructs the vector was digested with NdeI and EcoRI.

For constructs with the N terminal flag epitope, pXGFPN-5 was amplified in two steps to append the M2 epitope using 1) ORZ1871 and ORZ1763 2) ORZ1872 and ORZ1762.

For constructs with the C terminal flag epitope, pXGFPN-5 was amplified in two steps to append the M2 epitope using 1) ORZ1758 and ORZ1872 and 2) ORZ1871 and ORZ1759. All vectors were assemble using isothermal assembly with equal molar concentrations of insert and vector. pXGFPN-5_CcLon (S674A) was constructed by amplifying the coding sequence of Lon with ORZ1664 and ORZ1566.

pXGFPN-5_M2CcLon was constructed by amplifying the coding sequence of Lon with ORZ1855 and ORZ1854.

pXGFPN-5_CcLonM2 was constructed by amplifying the coding sequence of Lon with ORZ1856 and ORZ1853.

pXGFPN-5_LonH799K and pXGFPN-5_LonH799D were constructed by amplifying the coding sequence of Lon with ORZ1664 and ORZ1882 or ORZ1883, respectively.

pMCS-1_Lon(M2) was constructed by amplifying the pXGFPN-5_Lon(M2) vector with ORZ1888 and ORZ1890.

### Viability plating

For viability spot assays *Caulobacter* strains were grown overnight in PYE media supplemented with glucose. After overnight growth, cells were back diluted to OD_600_ 0.1 and outgrown to mid-exponential phase before being normalized to OD_600_ =0.1 and 10-fold serially diluted on to media lacking additional sugars or containing xylose (0.2%) for induction or glucose (0.2%) for no induction. Plates were allowed to air dry for 2 hours prior to plating. Serial dilutions were grown for 2 days at 30°C prior to imaging.

### *In vivo* protein stability and western blot analysis

Stains were grown overnight in PYE media. After overnight growth, cells were back diluted to OD_600_ =0.1 and grown to mid-exponential phase. Samples were taken prior to translation shutoff to monitor steady state protein levels. Protein synthesis was inhibited by addition of 30 µg/ml chloramphenicol. Cell samples were removed at the indicated time points, normalized by OD_600_ and immediately lysed in SDS lysis buffer and stored at -20°C prior to western blot analysis. Lysed samples were boiled at 95°C for 10 min and then centrifuged at 15,000 RPM to clear cell debris. After centrifugation, the supernatant was run on 10% SDS-PAGE gels. Proteins were transferred to a nitrocellulose membrane at 100V for 1 hour and probed with polyclonal rabbit anti-DnaA (1:5000 dilution), anti-CcrM (1:5,000), anti-Lon (1:2,000), anti-ClpP (1:10,000) and anti-ctrA (1:5,000) antibody. Proteins were visualized with goat anti-rabbit (1:5,000) using a Licor detection system (Licor Odyssey CLx). Final images were quantified using imageJ (20) and degradation rates were plotted using Prism.

### Microscopy

Phase contrast microscopy images are of exponentially growing cells (Zeiss AXIO ScopeA1) mounted on 1% PYE agar pads using a 100X objective. Cells were grown overnight in the presences of glucose. After overnight growth, cells were back diluted to OD_600_ 0.1 and grown to mid-exponential phase prior to imaging. Cell curvature was quantified using MicrobeJ (21) for ImageJ (20). Prism (Graphpad) was used for graphical representation of cell curvature measurements which is defined as the reciprocal of the radius of curvature. Representative images of the same scale were cropped to illustrate morphological defects.

### Protein Purification

DnaA was affinity purified by addition of His6SUMO to the N-terminus (22). His_6_SumoDnaA was expressed in Top10 *E. coli*. The cells were grown until OD_600_ (0.8), induced with 0.2% arabinose for 3 hours at 37°C shaking at 200 RPM. After induction cells were pelleted by centrifuged at 4°C 5,000 RPM for 10 mins. The pellets were resuspended in lysis buffer (20% sucrose, 20 mM HEPES (pH 7.5), 200 mM L-glutamic acid potassium, 10 mM MgCl_2_, 20 mM imidazole, and 1 mM DTT. Cell suspension containing 1 mM PMSF was lysed by microfluidizer (Microfluidics, Newton, MA). Lysate was cleared at 14,5000 RPM for 30 minutes at 4°C. Clarified lysate was immediately loaded onto a pre-equilibrated gravity Ni-NTA resin. The resin was washed three times with 3 column volumes lysis buffer and finally eluted with lysis buffer supplemented with 300 mM imidazole. The eluted protein was buffer exchanged into lysis buffer without imidazole and the His_6_SUMO tag was cleaved with Ulp1-his protease overnight at 4°C with gentle agitation. To further purify DnaA, cleavage products from Ulp protease cleavage were separated by additional ion-exchange column (GE healthcare, MonoS G5/50) using a KCl gradient of 0.1M to 1M in MonoS-Sucrose buffer (20% sucrose, 25 mM HEPES pH 7.5, 2 mM DTT). Native Lon, ClpA and ClpP protease was purified as previously described (23–26).

### In vitro degradation assay

Degradation for all reactions were performed at 30°C with the following protein concentrations unless elsewhere indicated: 0.1 μM Lon_6_ or 0.2 µM ClpAΔ9 and 0.4 µM ClpP, 2.5 μM DnaA, with 4 mM ATP, 15 mM creatine phosphate (Sigma) and 75 μg/mL creatine kinase (Roche) as ATP regeneration components. The reactions were initiated by adding the ATP regeneration mix to the protease-substrate solution in HEPES buffer (25 mM HEPES pH 7.5, 100 mM KCl, 10 mM MgCl_2_, and 1 mM DTT). Twenty μL aliquots were taken at each indicated time point and quenched in SDS loading dye (2% SDS, 6% Glycerol, 50 mM Tris pH 6.8 and 2 mM DTT), and examined by 10% SDS-PAGE gels.

## Acknowledgements

We thank the Chien, Strieter, Stratton, and Serio lab members for helpful comments and discussion on the manuscript. We thank Lucy Shapiro for providing CcrM antibody. This project was supported by funds from the NIH (R35GM130320 to P.C.). R.Z and S.M. were supported in part through the UMass NIH Chemistry Biology Interface Training Program (NIHT32GM008515). S.M. was funded in part by the HHMI Gilliam Fellowship Program. B.B was funded in part through the UMass Lee-SIP Undergraduate Summer Research Program.

---

**Figure S1.**
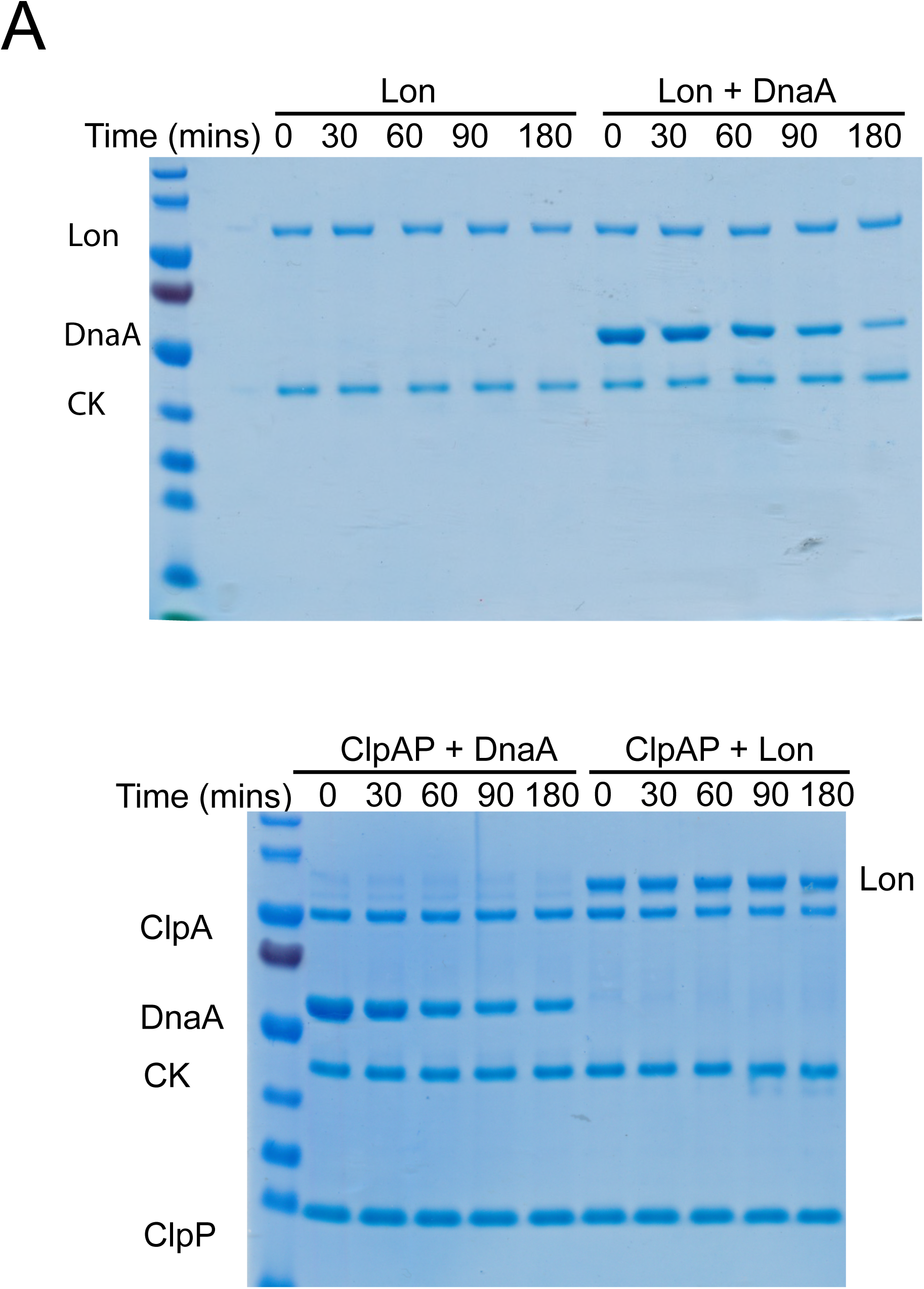
*in vitro* degradation assays with Lon and/or ClpAP. Degradation assays were performed using the native Lon substrate DnaA to test protease activity, Lon alone, ClpAP with Lon. Reactions consisted of (when listed) 2.5 µM DnaA, 0.1 µM Lon hexamer, 0.2 µM ClpA, 0.4 µM ClpP. All reactions contained 4 mM ATP and a suitable regeneration system (see methods)

## References

1. Sauer RT, Baker TA. 2011. AAA+ Proteases: ATP-Fueled Machines of Protein Destruction. Annu Rev Biochem 80:587–612.

2. Gottesman S. 2003. Proteolysis in Bacterial Regulatory Circuits. Annu Rev Cell Dev Biol 19:565–587.

3. Arsène F, Tomoyasu T, Bukau B. 2000. The heat shock response of Escherichia coli. International Journal of Food Microbiology 55:3–9.

4. Straus DB, Walter WA, Gross CA. 1987. The heat shock response of E. coli is regulated by changes in the concentration of s32. Nature 329:348–351.

5. Straus D, Walter W, Gross CA. 1990. DnaK, DnaJ, and GrpE heat shock proteins negatively regulate heat shock gene expression by controlling the synthesis and stability of sigma 32. Genes & Development 4:2202–2209.

6. Craig EA, Gross CA. 1991. Is hsp70 the cellular thermometer? Trends in Biochemical Sciences 16:135–140.

7. Bukau B. 1993. Regulation of the Escherichia coli heat-shock response. Mol Microbiol 9:671–680.

8. Jenal U. 1998. An essential protease involved in bacterial cell-cycle control. The EMBO Journal 17:5658–5669.

9. Christensen SK, Maenhaut-Michel G, Mine N, Gottesman S, Gerdes K, Van Melderen L. 2004. Overproduction of the Lon protease triggers inhibition of translation in Escherichia coli: involvement of the yefM-yoeB toxin-antitoxin system: Lon-dependent translation inhibition. Molecular Microbiology 51:1705–1717.

10. Ishii Y, Amano F. 2001. Regulation of SulA cleavage by Lon protease by the C-terminal amino acid of SulA, histidine. Biochem J 358:473–480.

11. Liu J, Francis LI, Jonas K, Laub MT, Chien P. 2016. ClpAP is an auxiliary protease for DnaA degradation in Caulobacter crescentus. Mol Microbiol 102:1075–1085.

12. Hottes AK, Shapiro L, McAdams HH. 2005. DnaA coordinates replication initiation and cell cycle transcription in Caulobacter crescentus: Caulobacter DnaA is a transcription factor. Molecular Microbiology 58:1340–1353.

13. Gottesman S, Clark WP, Maurizi MR. 1990. The ATP-dependent Clp protease of Escherichia coli. Sequence of clpA and identification of a Clp-specific substrate. J Biol Chem 265:7886–7893.

14. Seol JH, Yoo SJ, Kang M-S, Ha DB, Chung CH. 1995. The 65-kDa protein derived from the internal translational start site of the *clpA* gene blocks autodegradation of ClpA by the ATP-dependent protease Ti in *Escherichia coli*. FEBS Letters 377:41–43.

15. Gorbatyuk B, Marczynski GT. 2001. Physiological consequences of blocked Caulobacter crescentus dnaA expression, an essential DNA replication gene. Mol Microbiol 40:485–497.

16. Jonas K, Liu J, Chien P, Laub MT. 2013. Proteotoxic stress induces a cell-cycle arrest by stimulating Lon to degrade the replication initiator DnaA. Cell 154:623–636.

17. Wright R, Stephens C, Zweiger G, Shapiro L, Alley MR. 1996. Caulobacter Lon protease has a critical role in cell-cycle control of DNA methylation. Genes & Development 10:1532–1542.

18. Zeinert RD, Baniasadi H, Tu B, Chien P. 2019. The Lon protease links nucleotide metabolism with proteotoxic stress. bioRxiv 870733; doi: https://doi.org/10.1101/870733

19. Thanbichler M, Iniesta AA, Shapiro L. 2007. A comprehensive set of plasmids for vanillate- and xylose-inducible gene expression in Caulobacter crescentus. Nucleic Acids Res 35:e137–e137.

20. Schneider CA, Rasband WS, Eliceiri KW. 2012. NIH Image to ImageJ: 25 years of image analysis. Nat Methods 9:671–675.

21. Ducret A, Quardokus EM, Brun YV. 2016. MicrobeJ, a tool for high throughput bacterial cell detection and quantitative analysis. Nature Microbiology 1:16077.

22. Wang KH, Sauer RT, Baker TA. 2007. ClpS modulates but is not essential for bacterial N- end rule degradation. Genes Dev 21:403–408.

23. Gur E, Sauer RT. 2008. Recognition of misfolded proteins by Lon, a AAA+ protease. Genes Dev 22:2267–2277.

24. Levchenko I, Seidel M, Sauer RT, Baker TA. 2000. A Specificity-Enhancing Factor for the ClpXP Degradation Machine. Science 289:2354.

25. Liu J, Zeinert R, Francis L, Chien P. 2019. Lon recognition of the replication initiator DnaA requires a bipartite degron. Molecular Microbiology 111:176–186.

26. Maglica Ž, Striebel F, Weber-Ban E. 2008. An Intrinsic Degradation Tag on the ClpA C-Terminus Regulates the Balance of ClpAP Complexes with Different Substrate Specificity. Journal of Molecular Biology 384:503–511.

